# Selective neural coding of object, feature, and geometry spatial cues in humans

**DOI:** 10.1101/2021.04.28.441776

**Authors:** Stephen Ramanoël, Marion Durteste, Alice Bizeul, Anthony Ozier-Lafontaine, Marcia Bécu, José-Alain Sahel, Christophe Habas, Angelo Arleo

## Abstract

Orienting in space requires the processing and encoding of visual spatial cues. The dominant hypothesis about the brain structures mediating the coding of spatial cues stipulates the existence of a hippocampal-dependent system for the representation of geometry and a striatal-dependent system for the representation of landmarks. However, this dual-system hypothesis is based on paradigms that presented spatial cues conveying either conflicting or ambiguous spatial information and that amalgamated the concept of landmark into both discrete 3D objects and wall features. These confounded designs introduce difficulties in interpreting the spatial learning process. Here, we test the hypothesis of a complex interaction between the hippocampus and the striatum during landmark and geometry visual coding in humans. We also postulate that object-based and feature-based navigation are not equivalent instances of landmark-based navigation as currently considered in human spatial cognition. We examined the neural networks associated with geometry-, object-, and feature-based spatial navigation in an unbiased, two-choice behavioral paradigm using fMRI. We showed evidence of a synergistic interaction between hippocampal and striatal coding underlying flexible navigation behavior. The hippocampus was involved in all three types of cue-based navigation, whereas the striatum was more strongly recruited in the presence of geometric cues than object or feature cues. We also found that unique, specific neural signatures were associated with each spatial cue. Critically, object-based navigation elicited a widespread pattern of activity in temporal and occipital regions relative to feature-based navigation. These findings challenge and extend the current view of a dual, juxtaposed hippocampal-striatal system for visual spatial coding in humans. They also provide novel insights into the neural networks mediating object vs. feature spatial coding, suggesting a need to distinguish these two types of landmarks in the context of human navigation.

**Highlights:**

- Complex hippocampal-striatal interaction during visual spatial coding for flexible human navigation behavior.
- Distinct neural signatures associated with object-, feature-, and geometry-based navigation.
- Object- and feature-based navigation are not equivalent instances of landmark-based navigation.

## Introduction

The ability to navigate in space is fundamental to most daily activities, whether it be choosing the shortest route to work or meeting a friend in a familiar neighborhood. Despite its apparent simplicity, spatial navigation is a highly complex cognitive process that requires the integration of multimodal sensory information, the creation and maintenance of spatial representations in memory, and the manipulation of these representations to guide navigational behavior effectively (Wolbers & Hegarty, 2010). The adequate use of visual spatial cues constitutes an essential aspect of this process in many species, including humans (Ekstrom, 2015; Gouteux et al., 2001; Hermer & Spelke, 1996).

Early studies pointed to landmarks and geometry as distinct types of visual spatial cues used for self-orientation and navigation (Cheng, 1986; Gallistel, 1990). In the literature, the term *landmark* has been used to designate both discrete elements such as objects and buildings and embedded *featural* information such as color and texture (Doeller et al., 2008; Epstein & Vass, 2014; Gouteux & Spelke, 2001; Lowe et al., 2017; Marchette et al., 2015; Sutton et al., 2010; Wolbers & Büchel, 2005). The term *geometry* has been used to refer to the information provided by layouts such as relative lengths, distances, and angles between surfaces. While some argue that such information can only be derived from three-dimensional extended surfaces, others warrant a more comprehensive definition of geometry that includes the implicit relationships between objects (Cheng, 1986; Gouteux & Spelke, 2001; Learmonth et al., 2008; Lee & Spelke, 2010; Lourenco et al., 2009; Sutton et al., 2012).

The presence of visual spatial cues in an environment enables long-term spatial knowledge by facilitating the formation of cognitive maps (Epstein & Vass, 2014). Landmarks’ size, stability and proximity to the goal are among the key factors that determine their validity as anchor points and their use for navigation (Auger et al., 2015; Auger & Maguire, 2018; Stankiewicz & Kalia, 2007). Geometry constitutes a highly stable and indispensable information as it sets the environment boundaries, thus delineating the navigability of a space (Bécu et al., 2020; Keinath et al., 2017; Sheynikhovich et al., 2009). Critically, the availability, relative importance and complexity of visual spatial cues in the environment also influence the choice of navigation strategy (Arleo & Rondi-Reig, 2007; Foo et al., 2005; Keinath et al., 2017; O’Keefe & Nadel, 1978; Packard & Goodman, 2013; Ratliff & Newcombe, 2008). Two main navigation strategies can be employed: place strategies, which rely on flexible cognitive map-like representations of the environment; and response strategies, which rely on the formation of associations between a specific cue and a directional behavior (Arleo & Rondi-Reig, 2007; Kosaki et al., 2018). Given the intricacy of spatial navigation abilities in humans, it is perhaps not surprising that an extended neural network encompassing occipital, parietal, temporal and frontal regions is recruited (Epstein et al., 2017; Qiu et al., 2019). Decades of lesion studies and functional magnetic resonance imaging (fMRI) research have established the hippocampus and associated structures of the medial temporal lobe, such as the entorhinal, perirhinal and parahippocampal cortices, as central nodes of this network (Epstein et al., 2017; Julian et al., 2018; Spiers & Barry, 2015). The ability to navigate using a place strategy has largely been ascribed to the hippocampus while the striatum is known to support response strategies and habitual behavior (Chersi & Burgess, 2015; Goodroe et al., 2018).

Few experiments exploring the neural underpinnings of human spatial navigation have manipulated the presence of landmark and geometric cues in the environment. In an influential fMRI study by Doeller and colleagues in which participants had to learn locations relative to a salient object or to a circular enclosure, a striatal-dependent system for the representation of landmarks and a hippocampal-dependent system for the representation of geometry were uncovered (Doeller et al., 2008). This dual-system hypothesis forms the current widespread framework about the neural structures underlying landmark vs. geometry spatial coding. However, a limitation of this study lies in the complexity of the paradigm used. In their task both types of visual spatial cues were concomitantly present. Furthermore, geometric information consisted of a circular layout that could not be used alone as an orienting cue.

In a subsequent study, Sutton et al. (2010) conducted a fMRI experiment in which participants performed a navigation task within two separate virtual environments that each contained a single orienting cue (Sutton et al., 2010). The authors showed that reorientation based on a featural landmark (colored wall) elicited several activations within the medial temporal lobe whereas reorientation based on geometry (room shape) recruited the prefrontal and inferior temporal cortices. We argue however that the two environments did not allow for a clear dissociation of the neural circuits subtending the use of each cue subtype. Indeed, room size and cue reliance were not made equivalent in the landmark and geometry conditions. While reorienting with the colored wall was unambiguous, reorienting with geometry in the rectangular room was associated with a correct corner and an incorrect rotationally equivalent corner (Forloines et al., 2019; Sutton et al., 2012; Sutton & Newcombe, 2014). Moreover, it is worth noting that while Sutton and colleagues defined landmarks as featural information (colored wall), Doeller et al. defined landmarks as discrete objects independent of the boundaries (vase) (Bullens et al., 2010; Doeller et al., 2008; Doeller & Burgess, 2008). The concept of *landmark* thus seems to vary greatly across studies and deserves clarification.

The inconsistent results mentioned before could originate from the paradigms used (Doeller et al., 2008; Sutton et al., 2010), in which the visual complexity and the ambiguity of the cues (i.e., landmark and geometry provide equivalent spatial information) hindered the possibility to isolate neural circuits for the use of distinct visual spatial cues (Lew, 2011). An alternative non-exclusive explanation lies in the diversity of interpretations regarding the notion of landmark in the literature. The present fMRI study aimed at identifying the specific neural correlates that underlie the processing of objects, features and geometry using an unbiased reorientation navigation task in a virtual maze environment (Figure 1). Specifically, we explored two main questions. First, we tested the validity of the classic framework stating two causally dissociable systems in the hippocampus and striatum. Secondly, we posited that object-based and feature-based navigation would not be equivalent forms of landmark-based navigation in terms of behavioral and neural markers. We used the term *object* to refer to discrete landmarks that are independent of the environment’s boundaries and *feature* to refer to salient information embedded within the boundaries. The virtual environment consisted of a sparse and neutral layout, and the task design allowed for the three different types of visual spatial cues to be clearly separated. We conducted a whole-brain analysis to determine how cerebral regions would be similarly or differentially involved in object, geometry-, and feature-based navigation. Building on the work by Doeller and colleagues (2008) and the substantial literature on the neural substrates of place vs. response strategy (Goodroe et al., 2018; Kosaki et al., 2018; Packard & Goodman, 2013), univariate and multivariate ROI analyses were then performed to elucidate the respective roles of the hippocampus and striatum in cue-based navigation.

**Figure 1.**
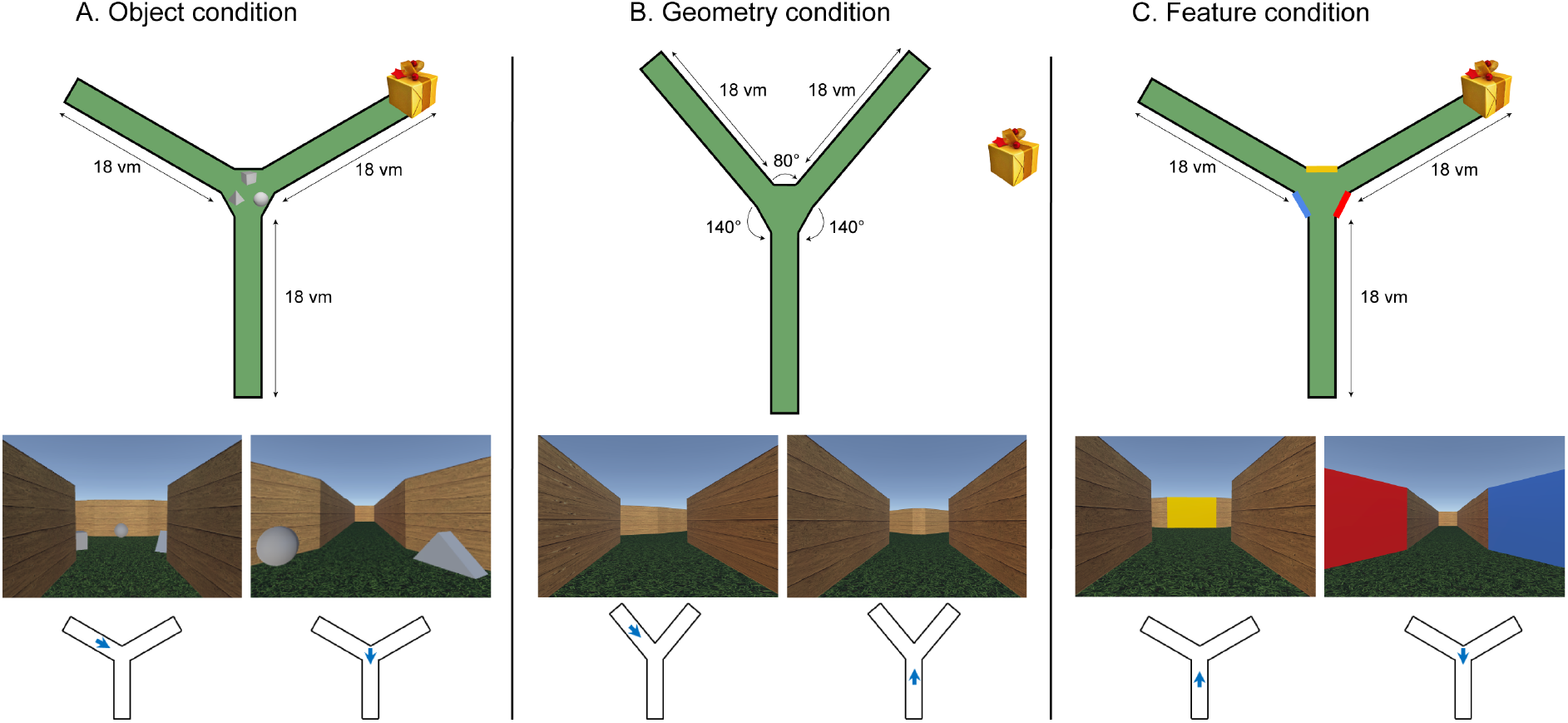
The virtual environment. **Top**: schematic overhead perspectives of the virtual environment for the object, geometry, and feature conditions. **(A)** In the object condition all arms were 18 virtual meters (vm) long and equiangular and three light gray objects were placed in the center of the maze. **(B)** In the geometry condition the arms were approximately 18 vm long, separated by two 140° angles and one 80° angle and there were no objects in the center of the maze. **(C)** In the feature condition, the arms were 18 vm long and equiangular, and there were three differently colored walls in the center of the maze. Of note, the overhead perspective was never seen by subjects. **Middle**: two sample first-person perspectives within the Y-maze for the object, geometry, and feature conditions. **Bottom**: blue arrows represent the position and direction associated with the first-person perspective views in each condition.

## Materials and Methods

### Participants

Twenty-six healthy young adults performed the fMRI experiment, but one participant was excluded due to poor task understanding. The final sample consisted of twenty-five participants (age 22-32, M = 25.4, SD = 2.7 years; 7F). Subjects were recruited from the French longitudinal cohort study *SilverSight* established in 2015 at the Vision Institute, Quinze-Vingts National Ophthalmology Hospital, Paris (Lagrené et al., 2019). Participants had no history of neurological or psychiatric disorders, they were right-handed, and they had normal or corrected-to-normal eyesight. Prior to the experimental session, subjects completed a battery of neuropsychological tests including the Mini-Mental State Examination (MMSE) (Folstein et al., 1975) and computerized versions of the 3D mental rotation test (Vandenberg & Kuse, 1978), the perspective-taking test (Kozhevnikov & Hegarty, 2001) and the Corsi block-tapping task (Corsi, 1973). Notably, the object and control conditions were also analyzed and included as control data in a study on healthy aging (Ramanoël et al., 2020).

All subjects provided written informed consent and the study was approved by the Ethical Committee “CPP Ile de France V” (ID_RCB 2015-A01094-45, CPP N°: 16122).

### Experimental design

#### Object, geometry, and feature definition

We defined objects as salient cues that are independent of the environment’s layout, geometry as the elements that are intrinsic to the external limits of a space (i.e., wall lengths and angle sizes) and features as salient information that is embedded within the environment’s layout (i.e., color). Worthy of note, the nature of geometric information can be separated into global geometry defined by relative wall lengths, and local geometry defined by angle dimensions (Hupbach & Nadel, 2005; Reichert & Kelly, 2011). As this is a topic of ongoing debate, we chose not to make the distinction.

#### The virtual navigation task

Participants performed a reorientation task in a virtual environment using an MRI-compatible ergonomic two-grip response device (NordicNeuroLab, Bergen, Norway). The task was projected on a MRI-compatible liquid crystal display monitor (NordicNeuroLab, Bergen, Norway) positioned at the head of the scanner bore. The virtual environment was designed with Unity 3D game engine.

Inside the scanner, participants navigated actively in a Y-maze that consisted of three corridors radiating out from a center and delimited by homogenous wooden-like walls (Figure 1–A–B–C). Forward speed of movement was set at 3 virtual meters (vm)/s and turning speed at 40°/s. Subjects completed three distinct reorientation conditions (object, geometry and feature) and one control condition. In the object condition (OBJ), all arms were 18 vm long and equiangular, and three light gray objects were placed in the center of the maze (Figure 1–A). In the geometry condition (GEO), the arms were approximately 18 vm long, separated by two 140° angles and one 80° angle and there were no objects in the center of the maze (Figure 1–B). In the feature condition (FEAT), the arms were 18 vm long and equiangular, and there were three differently colored walls in the center of the maze (Figure 1–C). In the control condition (CTRL), the arms were 18 vm long and equiangular and the maze didn’t contain any salient visual spatial cues.

The object, geometry, and feature conditions comprised an encoding phase and a retrieval phase. For the encoding phase, participants started from the center of the maze. They had to locate and then remember the position of a goal hidden at the end of one of the three arms (gifts and balloons). To that end, subjects had to use the visual spatial cues available at the intersection. The encoding phase lasted just under 3 minutes, following which the retrieval phase in the same environment began. In each trial of the retrieval phase, participants started at the end of an arm that didn’t contain the goal and had to retrieve the hidden goal. The starting positions across trials were pseudorandomized between subjects. Once they reached the correct goal location, the gifts and balloons appeared, followed by a fixation cross on a grey background before the start of the next trial. Subjects needed to complete eight trials. The presentation order of the object, geometry, and feature conditions along with the starting corridor in each trial were counterbalanced between subjects. The control condition was always performed after the three cue-based reorientation conditions, and it consisted of a 4-trial retrieval phase only. Subjects started from the end of an arm chosen randomly and moved to the center of the environment from where the goal was readily visible. They then navigated towards it. The control condition was designed to account for confounding factors such as motor and simple perceptual aspects of the task. For all conditions, we recorded trial duration and response device use during active navigation. The response device’s sole purpose was to allow participants to freely navigate in space.

A debriefing phase outside the scanner in which participants were probed on their use of visual spatial cues concluded the experimental session. There were two possible options: (i) “I used a single cue to reorient”, (ii) “I used two or more cues to reorient”. The answers served to assess participants’ propensity for response-based or place-based navigation strategies. We categorized subjects into response-based strategy users when they relied on a single visual spatial cue (Chrastil, 2013; Packard & Goodman, 2013) and place-based strategy users when they relied on two cues or more (Colombo et al., 2017; Gazova et al., 2013; Iaria et al., 2003; Iglói et al., 2010, 2015; Laczó et al., 2017).

### MRI acquisition and data preprocessing

Data were acquired using a 3-Tesla Siemens MAGNETOM Skyra whole-body MRI scanner (Siemens Medical Solutions, Erlangen, Germany) equipped with a 64-channel head coil at the Quinze-Vingts National Ophthalmology Hospital in Paris. T2*-weighted echo-planar imaging (EPI) sequences, optimized to minimize signal dropout in the medial temporal region were used for functional image acquisition (voxel size = 3 x 3 x 2 mm, TR/TE/flip angle = 2685 ms/30 ms/90°, interslice gap = 1 mm, slices = 48, matrix size = 74 x 74, FOV = 220 x 220 mm). Finally, a T1-weighted high-resolution three-dimensional image was obtained using an MPRAGE sequence (voxel size = 1 x 1 x 1.2 mm, TR/TE/IT/flip angle = 2300 ms/2.9 ms/900 ms/9°, matrix size = 256 x 240 x 176).

All subsequent fMRI data analyses were performed using SPM12 release 7487 (Wellcome Department of Imaging Neuroscience, London, UK) and ArtRepair toolbox implemented in MATLAB R2015 (Mathworks Inc., Natick, MA, USA). To ensure state equilibrium, the first five functional volumes from each run were removed. Slice-timing correction and spatial realignment were applied before correcting for motion-related artefacts with ArtRepair. Volumes displaying elevated global intensity (fluctuation > 1.3%) and movement exceeding 0.5 mm/TR were repaired using interpolation from adjacent scans. The next preprocessing steps included co-registration with the T1-weighted anatomical scans, normalization to the Montreal Neurological Institute (MNI) space, and spatial smoothing with an 8 mm full-width half maximum (FWHM) Gaussian kernel.

### Statistical Analyses

#### Behavioral analyses

Repeated-measures ANOVA and chi-square tests were used to compare navigation performance across conditions, including success rate, navigation time, and strategy use. To analyze the impact of sex on navigation time and strategy use, Mann-Whitney U-tests and Fisher’s exact tests were performed. The potential impact of trial number on navigation performance was explored using Spearman rank correlations and repeated-measures ANOVA for each of the three cue-based conditions. Finally, associations between navigation time and scores on various neuropsychological measures were computed using Spearman rank correlations with a statistical threshold adjusted for multiple comparisons.

#### Whole-brain analyses

The general linear model (GLM) was used for block design for statistical analysis of fMRI data (Friston et al., 1995). Eight trials of the retrieval phase in the object condition, eight trials of the retrieval phase in the geometry condition, eight trials of the retrieval phase in the feature condition, four trials of the control condition, and fixation times were modeled as regressors, constructed as box-car functions and convolved with the SPM hemodynamic response function (HRF). The encoding phases were not taken into account as the time required to find the goal for the first time differed greatly between participants (Figure S1). Navigation time, response device use during active navigation as well as six movement parameters derived from the realignment correction (three translations, three rotations) were entered in the design matrix as covariates. Time series for each voxel were high-pass filtered (1/128 Hz cut-off) to remove low-frequency noise and signal drift. FMRI contrasts [OBJ > CTRL], [GEO > CTRL] and [FEAT > CTRL] were fed into a group-level one sample t-test to compute average activation maps. In addition, direct comparisons between the cue-based conditions were performed. The statistical threshold was set at *p* = 0.001 uncorrected with a minimum cluster extent of k = 10 voxels.

We also conducted a conjunction analysis to map the brain regions that were similarly activated in all three cue-based conditions. Conjunction was performed using the intersection of supra-threshold voxels for the three separate one-sample t-tests using a minimal extended threshold set at 10 voxels. Using a similar approach, we then performed a conjunction analysis for each pair of visual spatial cue type.

#### Region-of-interest analyses

##### Univariate analyses

The hippocampus and striatum (caudate and putamen nuclei) were defined from the AAL probabilistic brain atlas (Tzourio-Mazoyer et al., 2002). Mean fMRI parameter estimates for the contrasts [OBJ > CTRL], [GEO > CTRL] and [FEAT > CTRL] were extracted from the hippocampus and the striatum using the REX MATLAB-based toolkit. A two-way ANOVA was then performed to study the main effects and interactions of ROI (hippocampus and striatum) and condition (object, geometry, feature) on fMRI parameter activity. fMRI parameter estimates in the striatum were compared between place and response strategy users for the contrasts [OBJ > CTRL] and [FEAT > CTRL]. To control for a putative learning effect during the retrieval phase, mean parameter estimates in the hippocampus and striatum were also extracted across trial numbers in the object condition using the following contrasts: [OBJ t1 > CTRL t1], [OBJ t2 > CTRL t2], [OBJ t3 > CTRL t3], [OBJ t4 > CTRL t4], [OBJ t5 > CTRL t4], [OBJ t6> CTRL t4], [OBJ t7 > CTRL t4] and [OBJ t8 > CTRL t4]. The same analyses were conducted for the geometry and feature conditions. In order to obtain a measure of the association between performance in the task and ROI activity, we conducted Spearman rank correlations between hippocampal and striatal activity and navigation time in each condition. Finally, based on several reports suggesting functional differences along the antero-posterior axis of the hippocampus (Dalton et al., 2019; Zeidman & Maguire, 2016), we conducted complementary analyses on the anterior hippocampus (aHC) and posterior hippocampus (pHC) delineated from the Human Brainnetome Atlas (Fan et al., 2016).

##### Multivariate analyses

We used representational similarity analysis to further characterize the roles of the hippocampus and striatum in discriminating between the three types of spatial cues. We first re-estimated the aforementioned GLM (see Whole-brain analysis) on unsmoothed data. The intensity of the MR signal was extracted from the two pre-defined ROIs and averaged across trials of each unique condition. For each subject, we then constructed representational similarity matrices (RSMs) by calculating pairwise Pearson correlations between each multi-voxel pattern of activity from the object, geometry, and feature conditions. We thus obtained symmetric 3 x 3 matrices for the hippocampus and striatum with off-diagonal values specifying the neural similarity between each pair of conditions. One-sample permutation tests were performed on the average RSMs to test whether the pairwise correlation coefficients were significantly different from zero. To explore the differences in pattern similarity between the hippocampus and the striatum we conducted a linear mixed-effects model. The lower triangular correlation coefficients of each RSM were fisher-z transformed and included as the dependent variable in the model with ROI, condition pair and their interaction as fixed effects and a random intercept for participants. In order to further test the existence of two dissociable systems in the HC and striatum for the processing of geometry and objects respectively, two theoretical RSMs were constructed. The object and geometry RSMs were created as binary RSMs based on whether one type of spatial cue would be preferentially processed in a ROI (1 was assigned to highly similar pairs of conditions and 0 to highly dissimilar pairs of conditions). Separate linear mixed-effects models were performed to compare the lower triangular correlation coefficients of the theoretical and neural RSMs. For each theoretical RSM, the neural similarity values were included as the dependent variable with the spatial cue similarities included as fixed effects and participants as random intercepts. All the above-mentioned analyses were conducted using the nltools and netneurotools packages and custom code in Python except for the linear mixed-effects models which were performed using the lme4 package in R.

### Data and code availability statement

The behavioral data and custom codes used in this manuscript are available upon request, without any restrictions. The anonymized neuroimaging data could be made available upon reasonable request.

## Results

### Behavioral results

Success rate (i.e., the number of times a participant chose the correct corridor across trials) was equivalent in the three cue-based conditions (Figure 2–A–B). We reported significant differences in navigation time (i.e., time to reach the goal averaged across trials) between conditions (F(2, 24) = 5.55, *p* = 0.023, partial η^2^ = 0.32, 95% CI [0.02, 0.52]). Tukey post-hoc tests revealed that young adults took significantly longer to find the goal in the geometry condition than in the object condition (*p* = 0.031). The type of navigational strategy used differed significantly between cue-based conditions (*χ*^2^ (2, N = 25) = 17.65, *p* < 0.001, Cramer’s V = 0.49, 95% CI [0.24, 0.70]). A significantly lower proportion of participants used a place strategy in the geometry condition than in the object (*χ*^2^ (1, N = 25) = 15.79, *p* < 0.001, ϕ = 0.79, 95% CI [0.50, 0.92]) and feature conditions (*χ*^2^ (1, N = 25) = 15.79, *p* < 0.001, ϕ = 0.79, 95% CI [0.50, 0.92]); (Figure 2–C). There was no significant difference in navigation time between response and place strategy users in the object condition (11.99 ± 0.19 vs. 11.97 ± 0.13; t (23) = 0.07, *p* = 0.94) and feature condition (11.96 ± 0.19 vs. 12.01 ± 0.14; t (23) = 0.82, *p* = 0.26). Considering that sex differences in spatial navigation ability have been reported in the literature (Boone et al., 2018; Nazareth et al., 2019), we next verified whether an effect of sex was present in our data. We showed that success rate, navigation time and strategy use were equivalent between men and women (Figure S2). There were no significant differences in navigation time by trial number across conditions lending support to the idea that participants displayed optimal performance from the first trial (Figure 2–D). We can nonetheless note that navigation time seemed to decrease across trials of the geometry condition and that this relationship trended to significance. Finally, we found no significant correlations between neuropsychological test scores and navigation time in any of the three cue-based conditions.

**Figure 2.**
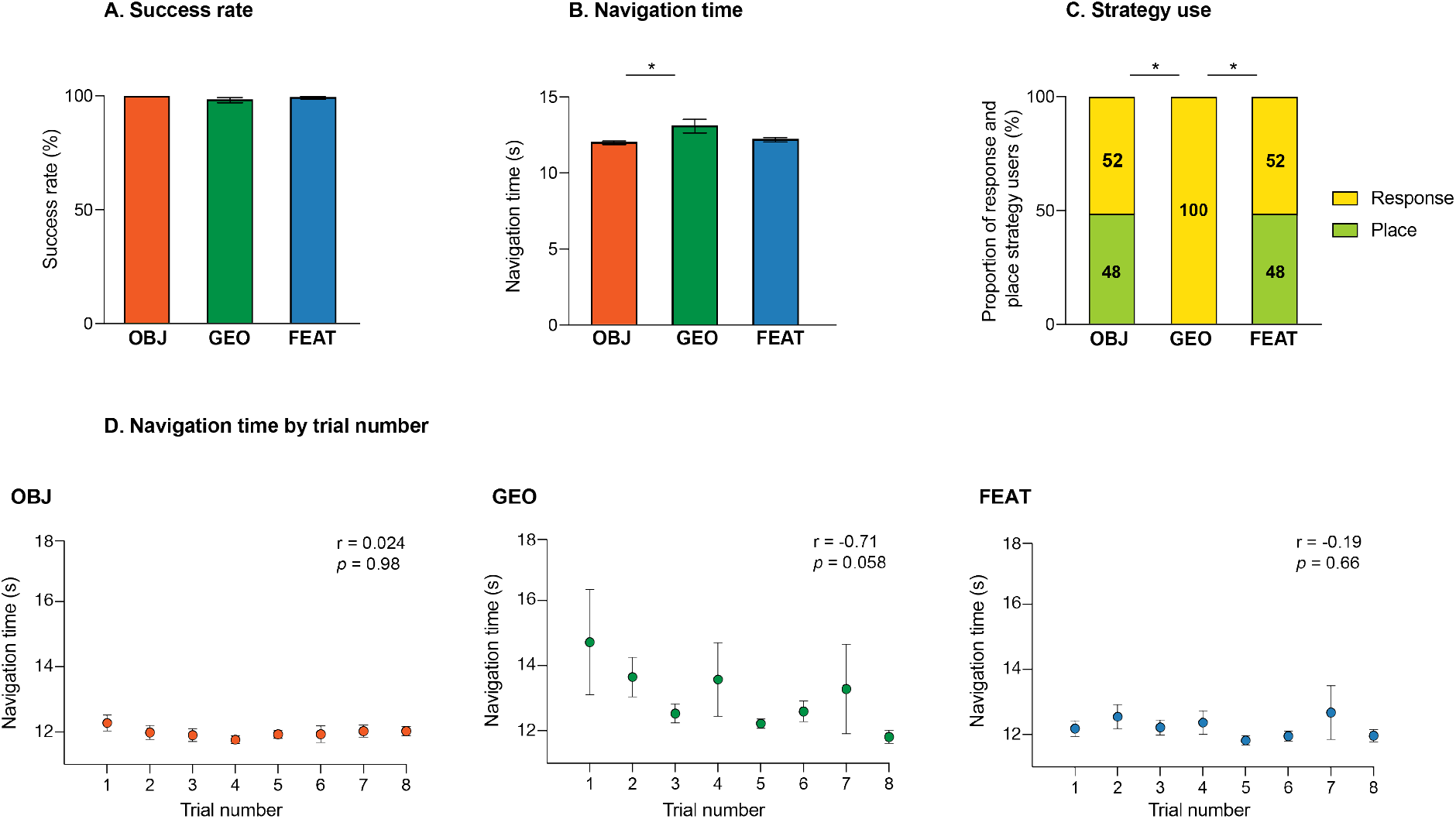
Behavioral results for the virtual navigation task across cue-based conditions. **(A)** Proportion of trials in which the correct corridor was chosen (success rate). **(B)** Time taken to reach the goal averaged across 8 trials (navigation time). **(C)** Proportion of participants using place-based or response-based strategies (strategy use). **(D)** Navigation time across trial number for the object condition (OBJ), geometry condition (GEO) and feature condition (FEAT). R and p-values correspond to Spearman rank correlations (top right). All errors bars represent standard errors of the mean.

### Whole-brain results

For all analyses, the navigation conditions were contrasted to the control condition using the fMRI contrasts [OBJ > CTRL], [GEO > CTRL] and [FEAT > CTRL]. We first conducted a conjunction analysis to explore the cerebral regions commonly activated by the three aforementioned contrasts (Table 1 and Figure S3). We reported significant activations in right anterior cingulate and right inferior temporal gyri. We then investigated the shared clusters of activation for each pair of visual spatial cues (Table 1). The object and geometry conditions both elicited activity in the fusiform gyrus bilaterally and in the right middle temporal gyrus while the object and feature conditions yielded activity in the right fusiform gyrus and in the right cerebellum. Of note, the conjunction analysis for the geometry and feature conditions showed no additional activation when compared to the main conjunction analysis that included the three conditions.

**Table 1.** Cerebral regions whose activity was elicited by the conjunction analyses between the three cue-based conditions contrasted to the control condition and between each pair of conditions contrasted to the control condition. The statistical threshold was defined as *p* < 0.001 uncorrected for multiple comparisons with an extent voxel threshold defined as 10 voxels. For each cluster, the region showing the maximum t-value was listed first, followed by the other regions in the cluster [in square brackets]. Montreal Neurological Institute (MNI) coordinates (x, y, z) of the peak and number of voxels (k) of clusters are also shown. OBJ = object condition; GEO = geometry condition; FEAT = feature condition; CTRL = control condition; H = hemisphere; R = right; L = left; BA = Brodmann area.

|  |  | H | BA | k | x | y | Z |
| --- | --- | --- | --- | --- | --- | --- | --- |
| **Conjunction Analysis** |  |  |  |  |  |  |  |
| **[OBJ ∩ GEO ∩ FEAT]** | Inferior Temporal Gyrus | R | 37 | 13 | 42 | -55 | -10 |
|  | Anterior Cingulate Gyrus | R | 24 | 11 | 6 | 34 | 4 |
| **[OBJ ∩ GEO]** |  |  |  |  |  |  |  |
| **[OBJ ∩ GEO]** | Fusiform Gyrus | L | 18 | 41 | -26 | -92 | -10 |
|  | Fusiform Gyrus | R | 18 | 39 | 33 | -87 | -8 |
|  | Anterior Cingulate Gyrus | R | 24 | 19 | 4 | 32 | 2 |
|  | Inferior Temporal Gyrus | R | 37 | 17 | 42 | -61 | -10 |
|  | Middle Temporal Gyrus | R | 37 | 11 | 43 | -55 | 10 |
| **[OBJ ∩ FEAT]** |  |  |  |  |  |  |  |
| **[OBJ ∩ FEAT]** | Inferior Temporal Gyrus | R | 37 | 27 | 41 | -60 | -11 |
|  | Fusiform Gyrus | R | 19 | 13 | 33 | -84 | -13 |
|  | Cerebellum | R |  | 11 | 30 | -55 | -43 |
|  | Anterior Cingulate Gyrus | R | 24 | 11 | 9 | 32 | 2 |
| **[GEO ∩ FEAT]** |  |  |  |  |  |  |  |
| **[GEO ∩ FEAT]** | Anterior Cingulate Gyrus | R | 24 | 35 | 9 | 32 | 4 |
|  | Inferior Temporal Gyrus | R | 37 | 16 | 42 | -64 | -10 |

Next, we were interested in differentiating the neural correlates associated with the use of objects, geometry, and features for spatial navigation. To this end, we looked at the fMRI contrasts [OBJ > CTRL], [GEO > CTRL] and [FEAT > CTRL] separately (Figure 3–A–B–C and Tables S2–S4). The object and geometry conditions ([OBJ > CTRL] and [GEO > CTRL]) elicited activity in bilateral superior temporal and right angular gyri. While the latter two conditions also elicited activation of the bilateral inferior occipital gyrus, the feature condition ([FEAT > CTRL]) yielded a significant cluster in the right inferior occipital gyrus. The geometry and feature conditions both elicited activity in the right superior frontal gyrus. Furthermore, we observed that left hippocampal and left inferior frontal activity were shared by both the object and feature conditions. Finally, multiple areas of the right cerebellum were found to be active in all three conditions.

**Figure 3.**
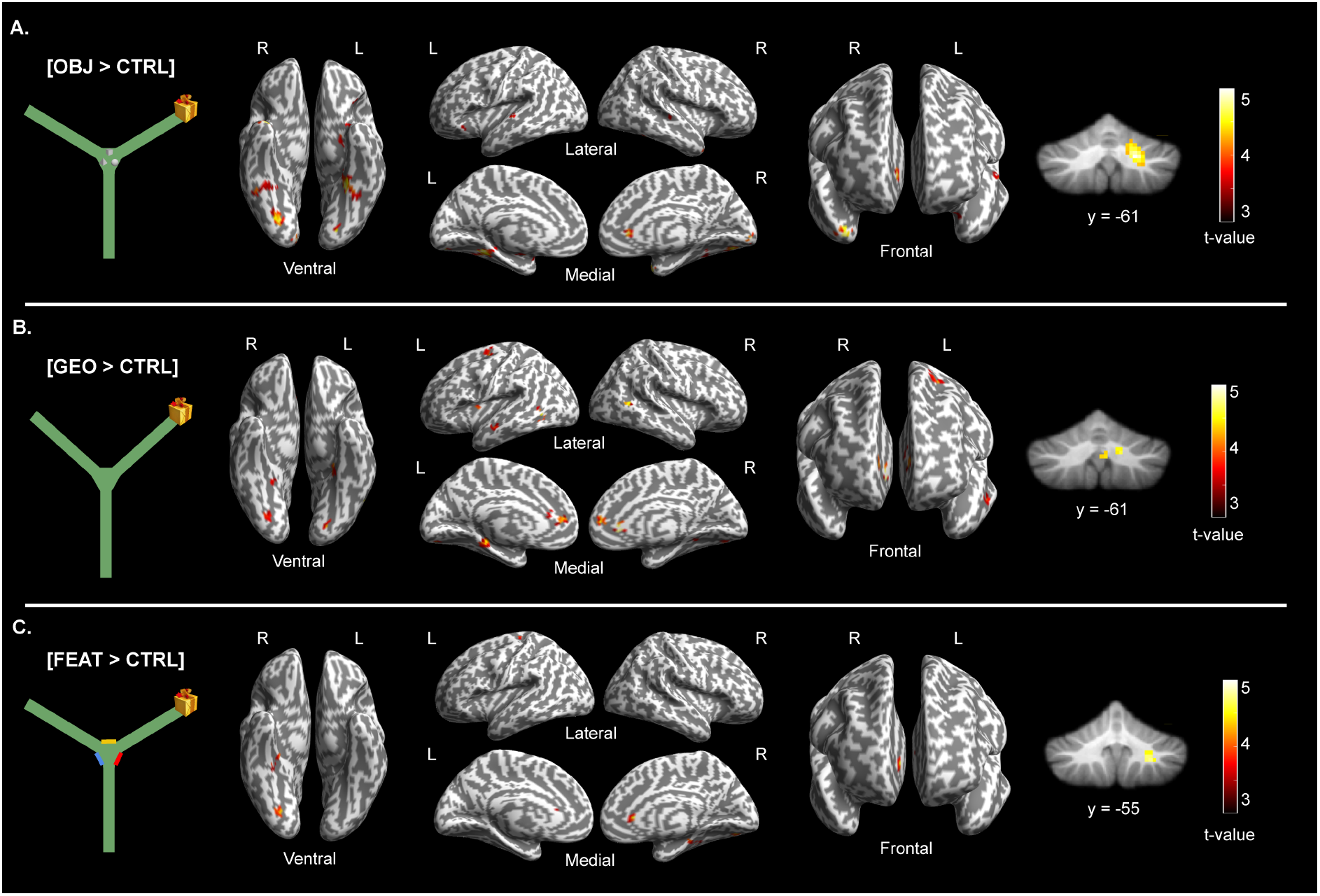
Cerebral regions whose activity was elicited by contrasting **(A)** the object condition with the control condition [OBJ > CTRL], **(B)** the geometry condition with the control condition [GEO > CTRL], and **(C)** the feature condition with the control condition [FEAT > CTRL]. The neural activity is projected onto 3D inflated anatomical templates and 2D slices for the cerebellum (*p* < 0.001 uncorrected, k = 10 voxels). OBJ = object condition; GEO = geometry condition; FEAT = feature condition; CTRL = control condition; L = left hemisphere; R = right hemisphere.

Examining the individual fMRI contrasts further revealed that each condition had specificities in its pattern of brain activity. The object condition elicited specific activations of bilateral middle occipital gyri and of the left lateral orbitofrontal gyrus. Moreover, the inferior temporal activity observed throughout the conditions comprised the anterior temporal pole only during object-based navigation. For the geometry condition, we noted extended activations in the frontal cortex encompassing clusters in the left superior frontal gyrus and left precentral gyrus. Geometry-dependent activations were also uncovered in the temporal lobes including the left parahippocampal gyrus and the left middle temporal gyrus, and in the right thalamus. The feature condition yielded activity in the left postcentral gyrus. Notably, the reverse fMRI contrasts, comparing the control condition with each navigation condition, did not reveal any significant activation (Tables S2–S4).

Finally, we conducted direct comparisons between the cue-based conditions themselves. Contrasting the geometry condition with the object condition ([GEO > OBJ]) elicited activity around the left superior frontal gyrus, extending to the left precentral gyrus (main cluster: -30, -10, 62), left caudate nucleus, left cerebellum and brainstem (Figure 4–A and Table S5). In addition, when contrasting the geometry condition with the feature condition ([GEO > FEAT]), we found significant activations in the bilateral middle temporal gyrus, middle frontal gyrus and putamen. We also reported activity in the left inferior occipital gyrus and left superior parietal gyrus as well as in the right supramarginal gyrus (Figure 4–B and Table S5). The fMRI contrasts comparing the object condition and feature condition with the other conditions ([OBJ > GEO], [OBJ > FEAT], [FEAT > OBJ], [FEAT > GEO]) did not elicit any significant activation (Table S5).

**Figure 4.**
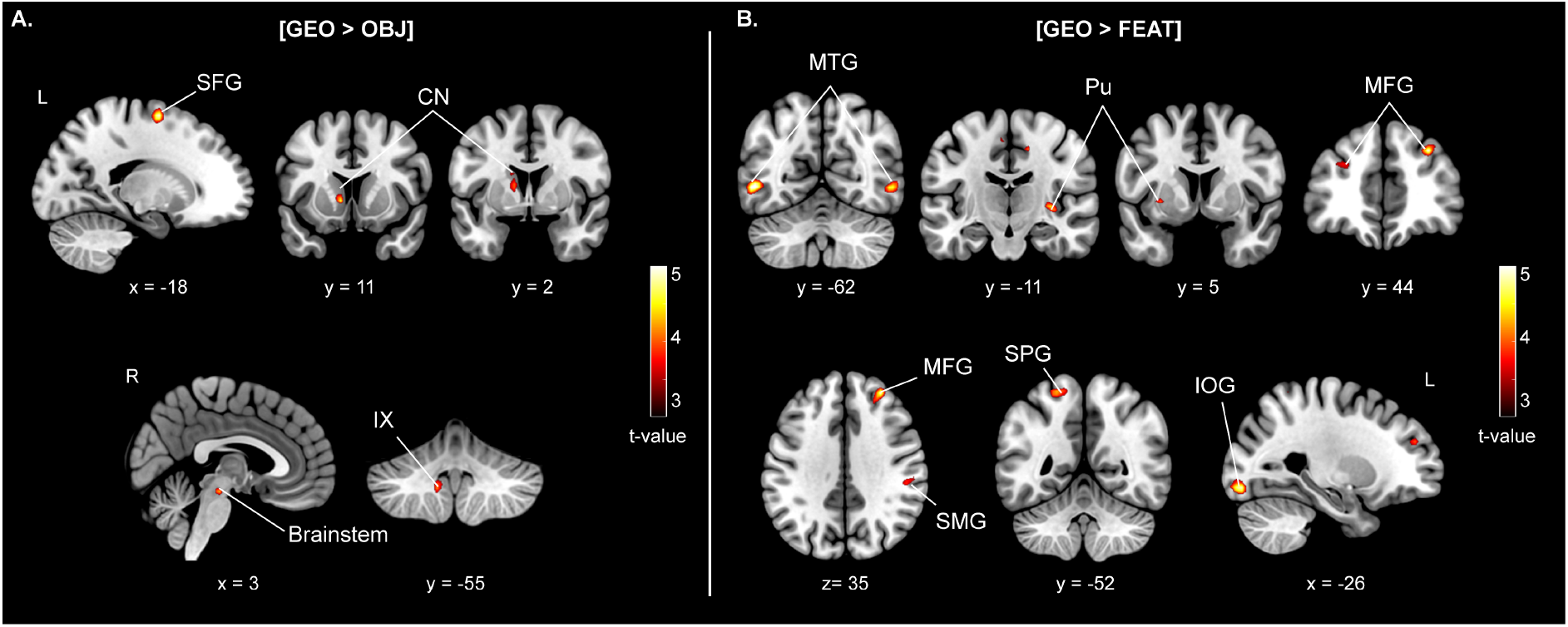
Cerebral regions whose activity was elicited by the fMRI contrasts **(A)** [GEO > OBJ] and **(B)** [GEO > FEAT]. The neural activity is projected onto 2D slices (*p* < 0.001 uncorrected, k = 10 voxels). OBJ = object; GEO = geometry; FEAT = feature; SFG = superior frontal gyrus; CN = caudate nucleus; IX = lobule IX cerebellum; MTG = middle temporal gyrus; Pu = putamen; MFG = middle frontal gyrus; SMG = supramarginal gyrus; SPG = superior parietal gyrus; IOG = inferior occipital gyrus; L = left hemisphere; R = right hemisphere.

### Regions of interest results

A repeated-measures two-way ANOVA was performed with ROI and condition as factors using the fMRI contrasts [OBJ > CTRL], [GEO > CTRL] and [FEAT > CTRL]. There were no main effects of ROI (F(1, 24) = 1.84, *p* = 0.19, partial η^2^ = 0.071, 95% CI [0.00, 0.30]) or condition (F(2, 48) = 1.56, *p* = 0.22, partial η^2^ = 0.061, 95% CI [0.00, 0.20) on parameter estimates (Figure 5–A). A significant interaction between ROI and condition was unveiled (F(2, 48) = 5.45, *p* = 0.007, partial η^2^ = 0.19, 95% CI [0.02, 0.35]. Post-hoc tests revealed that the hippocampus was more activated than the striatum in the object condition (0.40 ± 0.13 vs. -0.09 ± 0.11, *p* = 0.016, Hedge’s g = 0.48, 95% CI [0.06, 0.91]) and that there was decreased striatal activity in the object condition compared with the geometry condition (−0.09 ± 0.11 vs. 0.46 ± 0.22, *p* = 0.005, Hedge’s g = −0.54, 95% CI [−0.97, −0.12]).

**Figure 5.**
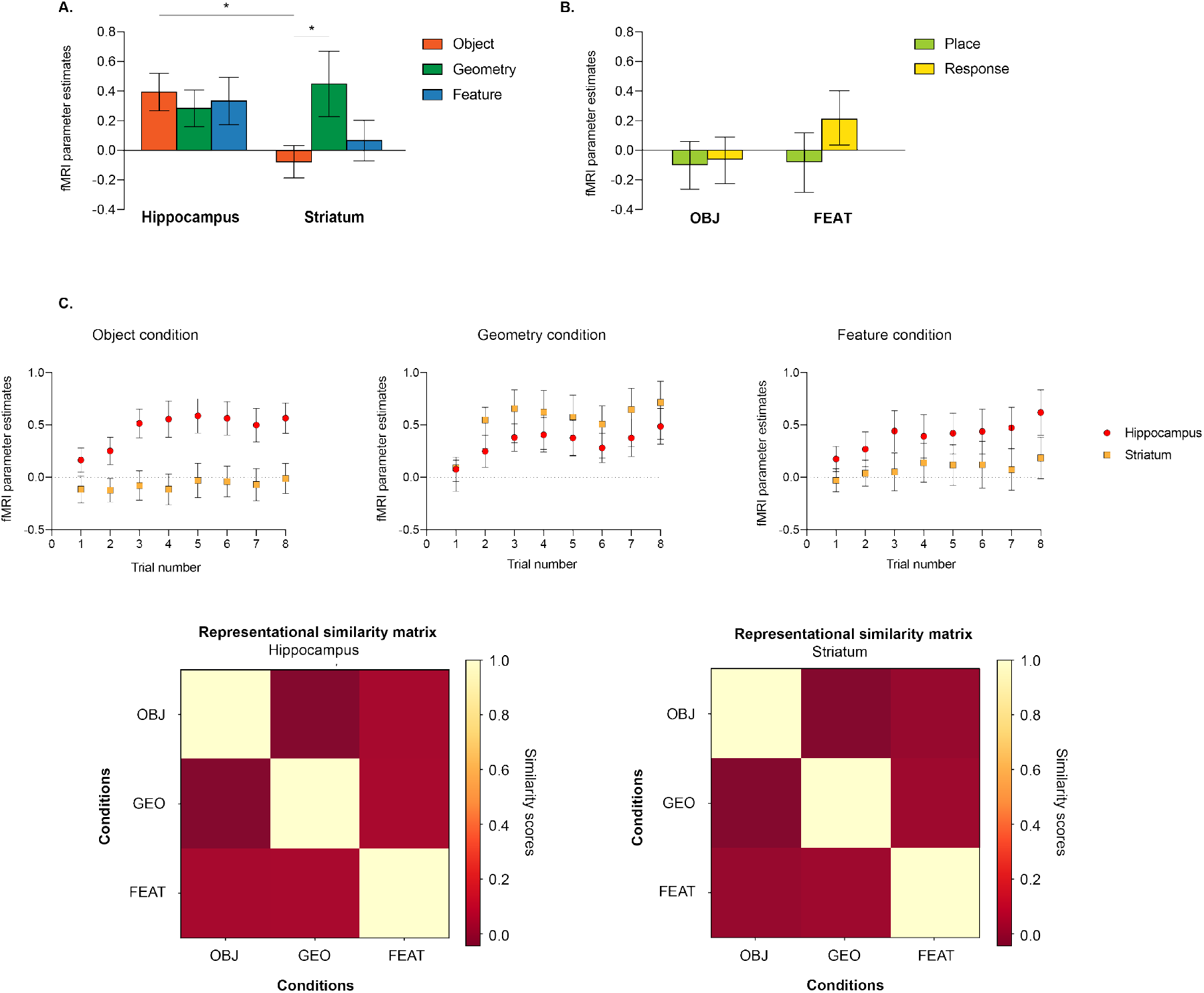
Results of the ROI analyses. **(A)** fMRI parameter estimates in the bilateral hippocampus and striatum for the fMRI contrasts [OBJ > CTRL], [GEO > CTRL] and [FEAT > CTRL]. **(B)** fMRI parameter estimates in the striatum for the fMRI contrasts [OBJ > CTRL] and [FEAT > CTRL] across place and response strategy users. **(C)** fMRI parameter estimates in the hippocampus and striatum for fMRI contrasts comparing neural activity in each trial of the object condition to that in one trial of the control condition (e.g., [OBJ t1 > CTRL t1], neural activity in each trial of the geometry condition to that in one trial of the control condition (e.g., [GEO t1 > CTRL t1]) and neural activity in each trial of the feature condition to that in one trial of the control condition (e.g., [FEAT t1> CTRL t1]. All error bars reflect standard errors of the mean. **(D)** Representational similarity matrices for the hippocampus (left) and striatum (right) averaged across subjects. Similarity scores are fisher z-transformed Pearson correlation coefficients, and they represent the overlap in terms of neural patterns between the object, geometry, and feature conditions

We observed the hippocampus to be implicated in all conditions, in varying degrees, and the striatum to be engaged solely during the geometry condition. Considering that the striatum is specifically implicated in response-based learning (Chersi & Burgess, 2015), we wondered whether the absence of striatal activity in the object and feature conditions could stem from differences in strategy use. We found that striatal activation was equivalent in the object and feature conditions between response and place strategy users (Figure 5–B). In order to identify putative learning effects, we examined fMRI parameter activity in individual trials for the contrasts [OBJ > CTRL], [GEO > CTRL] and [FEAT > CTRL]. There were no significant differences in hippocampal and striatal activity between trials for object, feature, or geometry conditions (Figure 5–C). We also conducted Spearman rank correlations between hippocampal and striatal activity and navigation time in each condition. We found a positive association between striatal activity and navigation time during geometry-based navigation only (r = 0.46, *p* = 0.014, 95% CI [0.10, 0.74]). Finally, we examined anterior and posterior hippocampal activity. These complementary analyses revealed that there were no significant differences in fMRI parameter activity across conditions between the anterior and posterior hippocampi (Figure S4).

To complement the above univariate findings, we conducted representational similarity analyses aimed at investigating the similarity of multi-voxel activations in the hippocampus and striatum between the object, geometry, and feature conditions. We found that the pairwise similarities between conditions were not significantly different from zero in the hippocampus and striatum (Figure 5–D). We then performed a linear mixed-effects model to test the influence of ROI, condition pair and their interaction (ROI x condition pair) on neural similarities. No significant effects were found. Separate linear mixed-effects model analyses for the hippocampus and striatum further revealed that neural similarity patterns were not significantly explained by neither the object (hippocampus: t = −1.49, *p* = 0.14; striatum: t = −0.84., *p* = 0.21) nor the geometry theoretical models (hippocampus: t = −1.44, *p* = 0.26; striatum: t = −0.31, *p* = 0.41).

## Discussion

Despite extensive knowledge surrounding the importance of visual information for spatial navigation, few studies have sought to elucidate how distinct types of visual spatial cues modulate behavior and brain activity. In this study, we examined the neural activity associated with object-, geometry-, and feature-based spatial navigation in an unbiased, two-choice behavioral paradigm using univariate and multivariate fMRI analyses. Success rate was equivalent across the three conditions. However, we found that participants took longer to reach the goal in the geometry condition than in the object condition and that the type of visual spatial cue available in the environment drove the use of different navigational strategies.

### A shared cerebral network for visual spatial cue processing

The whole-brain analyses revealed extended neural activations that were common to all conditions and that have been repeatedly implicated in spatial orientation paradigms (Chrastil, 2013; Cona & Scarpazza, 2019; Epstein et al., 2017; Qiu et al., 2019; Spiers & Barry, 2015). Unsurprisingly, multiple areas associated with visuospatial processing in the temporal and occipital lobes were activated. For example, the three conditions yielded activity in the fusiform gyrus, which may be reflecting the accurate recognition of novel information (Ewbank et al., 2005) and exemplifies the overarching importance of visual processing during fMRI virtual navigation tasks (Taube et al., 2013). We also noticed that the cerebellum was consistently activated, offering additional insight into its relevance for navigation (Iglói et al., 2015; Rochefort et al., 2013; Rondi-Reig et al., 2014).

While the above whole-brain results give us an idea of broad network similarities, the conjunction analysis revealed a more specific overlap of the right anterior cingulate cortex (ACC) and the right inferior temporal cortex across conditions. The common engagement of the ACC can be interpreted in light of its function as an internal monitor (Carter et al., 1998; Walton et al., 2003; C. Wang et al., 2005). Indeed, the primate ACC is positioned at the crossroads between medial temporal structures and premotor regions allowing for the integration of affective and contextual memory information with goal-directed action (Botvinick et al., 2004; Hayden & Platt, 2010; Ito et al., 2003). In our experiment, reorientation required participants to notice their change of position within the maze and re-evaluate their trajectory accordingly. Lending credence to our result, Javadi and colleagues (2019) similarly found peak ACC activation when subjects understood that they had deviated from the optimal path and that they needed to backtrack (Javadi et al., 2019). The inferior temporal cortex is a central part of the ventral visual stream and it is critical for high-level visual processing such as perceptual detection and identification of faces, objects and scenes (Aguirre et al., 1998; Epstein et al., 1999; Grill-Spector & Weiner, 2014; Kanwisher, 2010; Kravitz et al., 2013; Landis et al., 1986; Litman et al., 2009). Together, these findings strengthen the argument for similarities between distinct forms of visual spatial cue processing for navigation.

### The specificity of processing geometric spatial cues

Although participants displayed optimal performance in all three conditions in terms of success rate (Figure 2–A), our results emphasize a behavioral specificity for the processing of geometric cues compared with object and featural cues. Indeed, participants took longer to reach the goal in the geometry condition compared with the object condition and appeared to need more trials to reach optimal performance (Figure 2–B–D). Most strikingly, we found that all subjects used a response-based strategy to reorient with geometry while there were both place-based and response-based strategy users to reorient with objects or features (Figure 2–C). It is widely documented that the sensory cues available in the environment modulate behavior (Bosco et al., 2008; Foo et al., 2005; Kelly et al., 2009), but research investigating a specific place or response strategy bias is scarce.

Several important points can be made. First, we must stress that our definitions of place and response strategies were based solely on the number of cues participants used to orient. One can appreciate that the concept of geometry is less accessible in declarative memory than the concept of landmarks. In other words, it may have been easier for participants to be consciously aware of which visual spatial cues they were relying on in the object and feature conditions than in the geometry condition. Second, geometric information including angles between arms, lengths of corridors, and overall shape of the central area was available in the maze for participants to exploit. We speculate that this information was less noticeable and salient than both objects and features. The increased perceived difficulty of the geometry condition may provide an explanation for the differences in navigation time, learning curves as well as the overarching bias for response-based strategies. The questionnaire from the debriefing phase must therefore be apprehended cautiously and further investigation is warranted to gain a clearer understanding of the relationship between visual spatial cue type and navigational strategy.

The above behavioral results hint at the differential processing of geometric cues specifically and are corroborated by our fMRI whole-brain analyses. Indeed, our results highlight regional specificities for landmark- and geometry-based navigation. First, the whole-brain analyses revealed left hippocampal activity in the object and feature conditions only. It is important to emphasize that these results do not indicate an absence of hippocampal activity from the geometry condition, but rather that it is not significant compared to that in the other two conditions (see ROI results). Sutton and colleagues (2010) similarly reported increased hippocampal engagement in the presence of a distinctive featural cue embedded within environmental boundaries. We hypothesize that the saliency of object and featural cues enabled participants to better integrate local visual information with the broader spatial context into a unified hippocampal-dependent representation. Second, directly contrasting the geometry condition with the object and feature conditions revealed striking disparities. In accordance with previous studies, geometry-based navigation yielded greater activity in a vast neural network comprising the frontal cortex, parietal cortex and striatum (Forloines et al., 2019; Sutton et al., 2010, 2012). Both frontal and striatal regions are involved in the evaluation and selection of adequate behavior, and prefrontal inputs to the striatum could facilitate context-dependent and flexible navigational responses (Brown et al., 2012; Goodroe et al., 2018; Haber et al., 2006). Moreover, our ROI analyses revealed a positive correlation between navigation time and striatal activity in the geometry condition only. The specific demand on frontal and striatal function for geometry processing may thus indicate more intricate decision-making processes attributable to the reduced saliency of geometric cues. The greater parietal activity in the geometry condition compared with the object and feature conditions resonates with the exclusive use of a response-based strategy by participants when encoding geometric cues (Gauthier & van Wassenhove, 2016; Sherrill et al., 2015; Spiers & Maguire, 2007). Taken together, the above findings converge towards the idea that geometric cues in our paradigm are processed in a different manner to object and featural cues.

### Rethinking the concept of landmark in human spatial navigation

In fMRI studies, landmarks have been conceptualized as discrete objects (Doeller et al., 2008; Iaria et al., 2003; Janzen et al., 2008; Janzen & Van Turennout, 2004), abstract shapes (Baumann et al., 2010; Wegman et al., 2014), buildings (Wolbers & Büchel, 2005) and embedded visual information (Sutton et al., 2010). As previously emphasized by Mitchell and colleagues (2018), a pressing question remains as to how objects and features are considered by the brain and whether the field is correct in assuming their indistinguishability (Mitchell et al., 2018). Even though directly comparing the two conditions did not yield any significant activations, intriguing differences were noted when contrasting the two cue-based conditions individually to the control condition. First, object-based and feature-based navigation recruited distinct areas of the prefrontal cortex: the orbitofrontal gyrus and the superior frontal gyrus, respectively. While the orbitofrontal cortex is implicated in short-term memory for objects and is thought to mediate top-down visual recognition in association with temporal visual areas (Bar et al., 2006; Rolls, 2004), the superior frontal gyrus is more concerned with spatial working memory processing (Boisgueheneuc et al., 2006). The continuity of salient visual information with the environmental layout may thus activate regions oriented towards spatial processing rather than object processing. Second, we reported a more widespread pattern of activity comprising temporal and occipital regions during object-based navigation than during feature-based navigation. One can speculate that the integration of objects for spatial navigation requires fine-grained visual processing that is unnecessary to the use of salient colored walls (i.e., features). Along those same lines, we showed activation of the anterior temporal pole when contrasting the object condition with the control condition. This result fits with the gradient nature of the medial temporal cortex with the most anterior part being concerned with the processing of objects and the most posterior part with that of scenes (Kravitz et al., 2013). It is worth noting that activation of the postcentral gyrus was observed during the feature condition and not the object condition. Rarely reported in spatial navigation studies, this structure has once been proposed to play a specific role in the processing of spatial layout and local environment cue from a self-centered perspective (Shelton & Gabrieli, 2002). Further accentuating the dichotomy between objects and features, representational similarity analyses revealed no significant correlations between the neural patterns subtending the two conditions in the hippocampus and striatum. Thus, despite objects and features bearing equivalent permanence and spatial utility, higher cognitive structures of the brain appear to treat them differently. What is typically considered to be a landmark may influence the underlying patterns of neural activity. Indeed, one could argue that objects and colored walls convey very different types of information in the real-world with the former being less stable and more frequently interacted with than the latter. Finally, the distinct occipital activations also underline the possibility that visual properties such as angular size or color contributed to the observed neural differences between objects and features. These results have important implications. We revealed that while objects and features can be used equally well to orient in space, object-based navigation is subtended by a more widespread pattern of temporal and occipital activations. One could thus speculate that featural information demands fewer cognitive resources for efficient spatial navigation. Future research should test whether favoring featural instead of object cues in architectural designs could facilitate spatial orientation in complex indoor spaces.

### Synergistic interaction between the hippocampus and striatum

In light of the classic theory stating the existence of a hippocampal-dependent system for the representation of geometry and a striatal-dependent system for the representation of landmarks (Doeller et al., 2008; Doeller & Burgess, 2008), we conducted univariate and multivariate ROI analyses in the hippocampus and striatum. Interesting patterns emerged that we cautiously discuss below with a view to encourage further research. We observed equivalent ROI hippocampal activity in all three cue-based conditions and more striatal activity in the geometry condition. Moreover, representational similarity analyses revealed that the patterns of activity in the hippocampus and striatum weren’t specifically associated with the processing of either objects or geometry. Our results thus fail to support Doeller and colleagues’ conclusions that two causally dissociable systems (hippocampus-based and striatum-based) are associated with geometric and landmark cue processing respectively. Multiple factors could account for such discrepant findings. First, in the virtual environment designed by Doeller et al. both proximal and distal landmarks were available whereas our environment only contained proximal landmarks. Several reports have highlighted the behavioral and neural differences linked to processing proximal or distal visual information when navigating (Hébert et al., 2017; Knierim & Hamilton, 2011). Second, we used highly divergent definitions of *geometric information*. In their experiment geometry was a circular boundary that could not be used in itself to orient while geometry in our experiment consisted of angles and wall lengths and could be used to orient. The openness of the environmental space also constitutes a plausible candidate to explain our results. Multiple studies have shown that barriers modify spatial navigation performance and fragment the representation of space (Han & Becker, 2014; He et al., 2019; Li & Klippel, 2016; Meilinger et al., 2016; Wang & Brockmole, 2003). Doeller and colleagues’ environment consisted of a large open field whereas our virtual maze was a closed space delimited by corridors. Finally, we advocate that these contradictory findings can be best understood by discarding the view that the human hippocampus and striatum are largely juxtaposed systems during spatial cue encoding for spatial navigation. There is long-standing knowledge from rodent studies that the hippocampus mediates the formation of cognitive map-like representations (i.e., place-based strategies) while the striatum is specifically involved with stimulus-outcome associations (i.e., response-based strategies) (Squire, 2004; White & McDonald, 2002). Yet, converging evidence from the animal and human literature supports intricate cooperation and competition between these two memory systems (Gahnstrom & Spiers, 2020; Packard & Goodman, 2013; Rinaldi et al., 2020; van de Ven et al., 2020). Critically, Packard and Goodman (2013) posited that the heterogeneity of the visual environment, defined by the presence of multiple visual cues, could modulate this competition by favoring one type of strategy over another, ultimately diminishing competitive interference. In our experiment, the homogeneity of the geometry condition may have enhanced response-based learning by preventing some competition from the hippocampus, which could account for the presence of striatal activity. The latter may also provide an explanation for the longer navigation time observed during geometry-based navigation. On the other hand, the heterogeneity of the environments in the object and feature conditions may have ceased most competitive interference from the striatum.

### Conclusion and perspectives

Taken together, our results suggest that despite some shared activations in the inferior temporal and anterior cingulate gyri, each instance of cue-based navigation displays its specific neural signature and is subtended by complex hippocampo-striatal interactions. Gaining a deeper understanding of the relationship between the hippocampus and the striatum during spatial navigation could provide a more definitive answer regarding their involvement in landmark vs. geometry processing (Doeller et al., 2008; Doeller & Burgess, 2008; Schuck et al., 2015). Moreover, the divergence between object and feature spatial coding stresses the importance of considering vision and spatial navigation as tightly interwoven systems and the need to reevaluate the concept of landmark in the field of human spatial navigation (Meister & Buffalo, 2016, 2018; Nau et al., 2018).

Multiple questions still remain regarding the complex interaction between the visual environment, the navigational strategy choice and the neural processing of visual spatial cues. Cognitive properties such as permanence and spatial utility were equivalent across conditions hinting at the possible contribution of lower-level processes (Mitchell et al., 2018). While the influence of visual properties on brain activity in sensory and navigationally-relevant regions such as the hippocampus is being thoroughly investigated in the context of natural scene perception (Aly et al., 2013; Dima et al., 2018; Ghodrati et al., 2016; Groen et al., 2013, 2017; Kaiser et al., 2019; Kauffmann et al., 2015; Seijdel et al., 2020), it has seldom been explored during active spatial navigation. Spatial frequency content, contrast amplitude, angular size, and position in the visual field may constitute interesting research avenues. Recent studies have revealed that age-related impairments in navigational abilities could be partially explained by the decline of information processing in early visual regions (Koch et al., 2020; Ramanoël et al., 2020). Therefore, specific modulations of these basic visual properties could stabilize or even improve navigation performance. To test such hypotheses, eye-tracking methods coupled to neuroimaging constitute promising tools to test whether oculomotor behavior changes as a function of visual spatial cue quality and type and can predict navigational strategy use (Nau et al., 2018, 2020).

## Acknowledgments

The authors would like to express their gratitude to the men and women who took part in this study. We thank the Quinze-Vingts Hospital for allowing us to acquire the MRI data. We thank Konogan Baranton and Isabelle Poulain (Essilor International) for the manufacturing of MRI-compatible glasses.

## Author Contributions

Study design: SR, AB, AOL, MB, JAS, CH, AA; Data acquisition: SR, AB; Data processing: SR, MD, AB; Manuscript writing: SR, MD, MB, AA.

## Declaration of Interests

The authors declare that the research was conducted in the absence of any commercial or financial relationships that could be construed as a potential conflict of interest.

## Funding

This research was supported by the Chair SILVERSIGHT Agence Nationale de la Recherche (ANR-18-CHIN-0002), the LabEx LIFESENSES (ANR-10-LABX-65), the IHU FOReSIGHT (ANR-18-IAHU-01) and the Fondation pour la Recherche sur Alzheimer (FRA).

## Supplemental information

**Figure S1.**
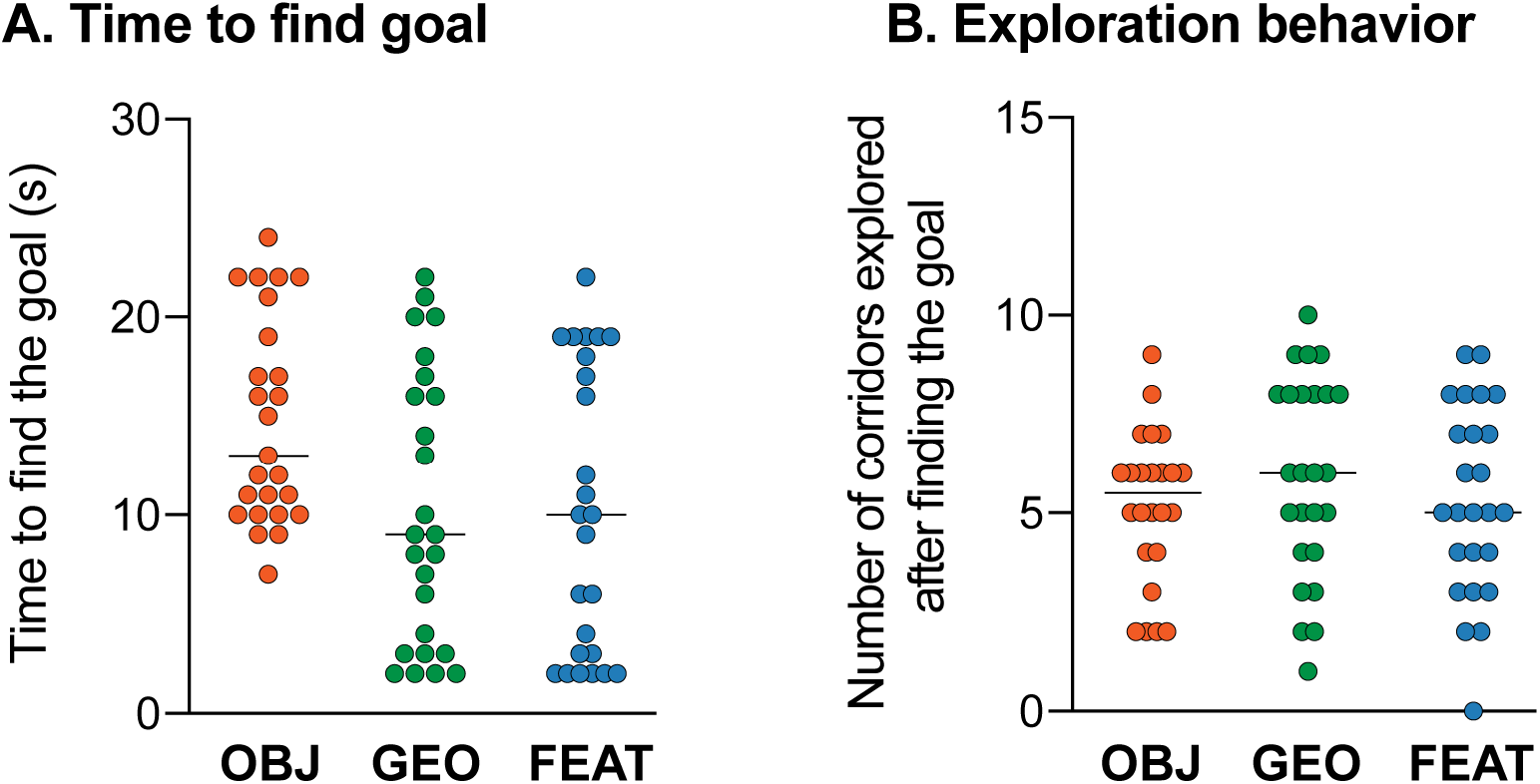
Behavioral results for the encoding phase of the virtual navigation task across cue-based conditions. **(A)** Time taken to find the goal for the first time for each participant. **(B)** Number of corridors explored by each participant after finding the goal. The horizontal black lines represent the median.

**Figure S2.**
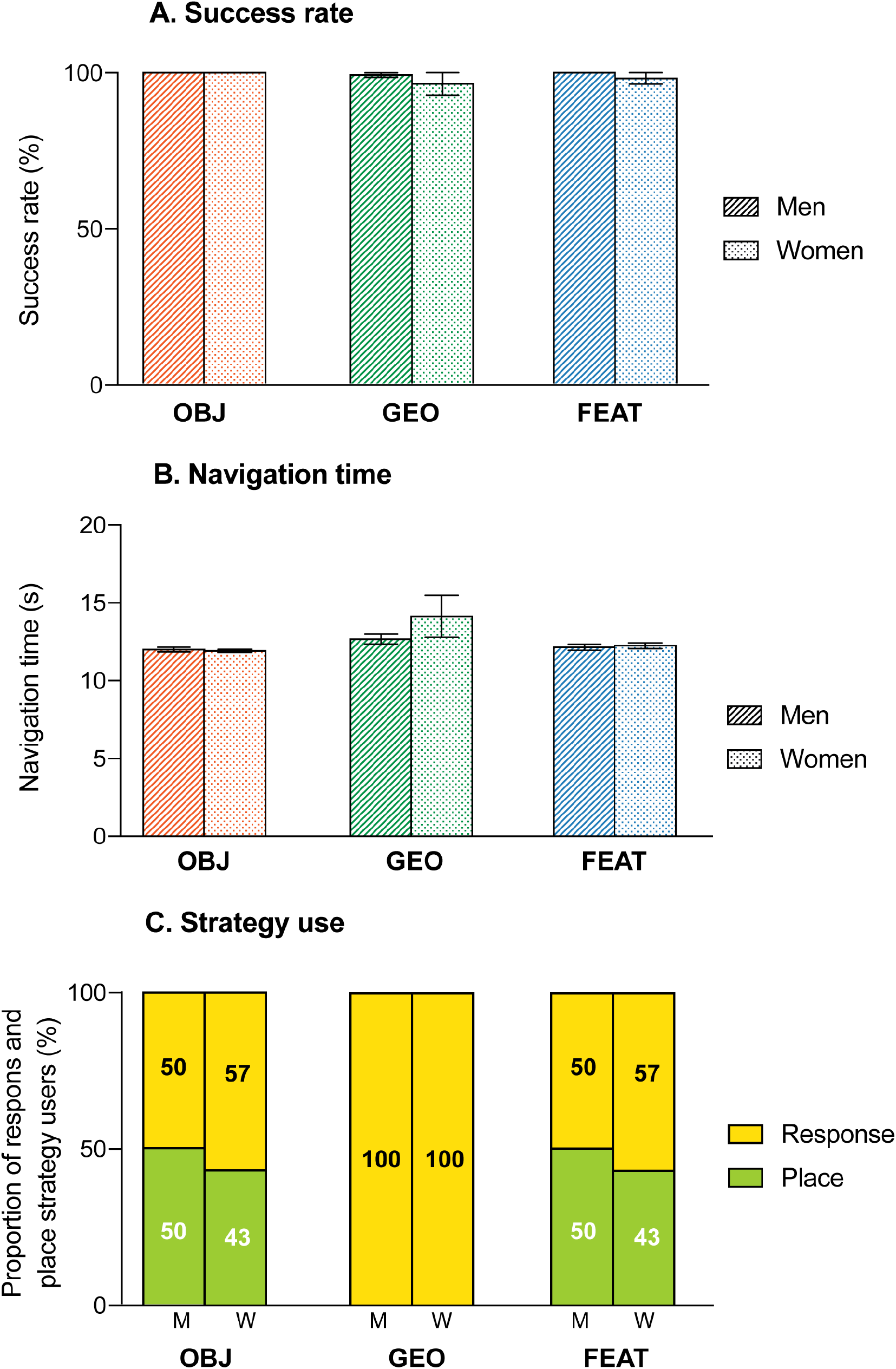
Behavioral results for the virtual navigation task across cue-based conditions in men and women separately. **(A)** Proportion of trials in which the correct corridor was chosen (success rate). **(B)** Time taken to reach the goal averaged across 8 trials (navigation time). **(C)** Proportion of participants using place-based or response-based strategies (strategy use). M = men; W = women. Error bars represent the standard errors of the mean.

**Figure S3.**
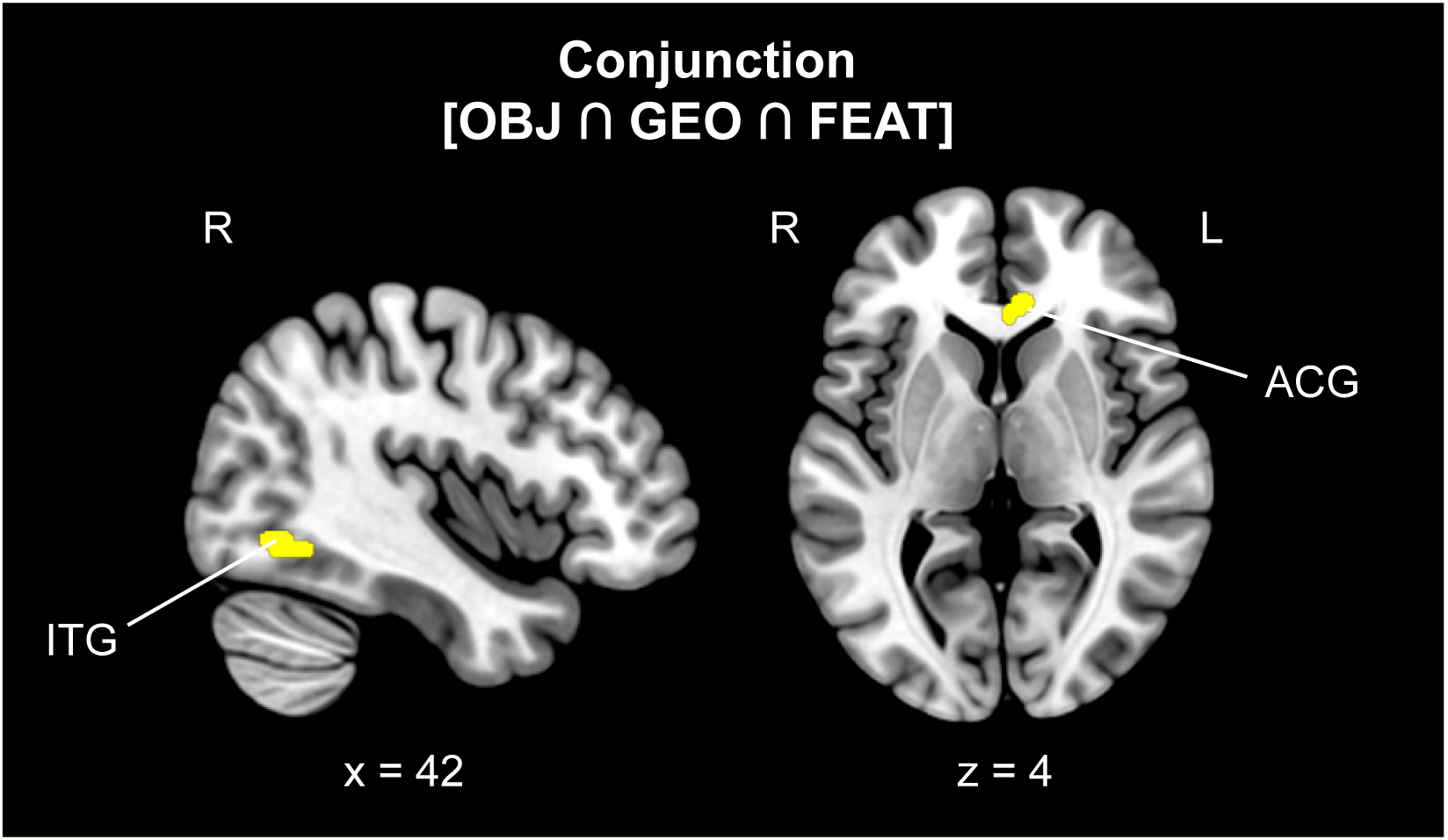
Cerebral regions whose activity was elicited by the conjunction analysis between the three cue-based conditions contrasted to the control condition. The neural activity is projected onto 2D slices (*p* < 0.001 uncorrected, k = 10 voxels). OBJ = object; GEO = geometry; FEAT = feature; ITG = inferior temporal gyrus; ACG = anterior cingulate gyrus; L = left hemisphere; R = right hemisphere.

**Figure S4.**
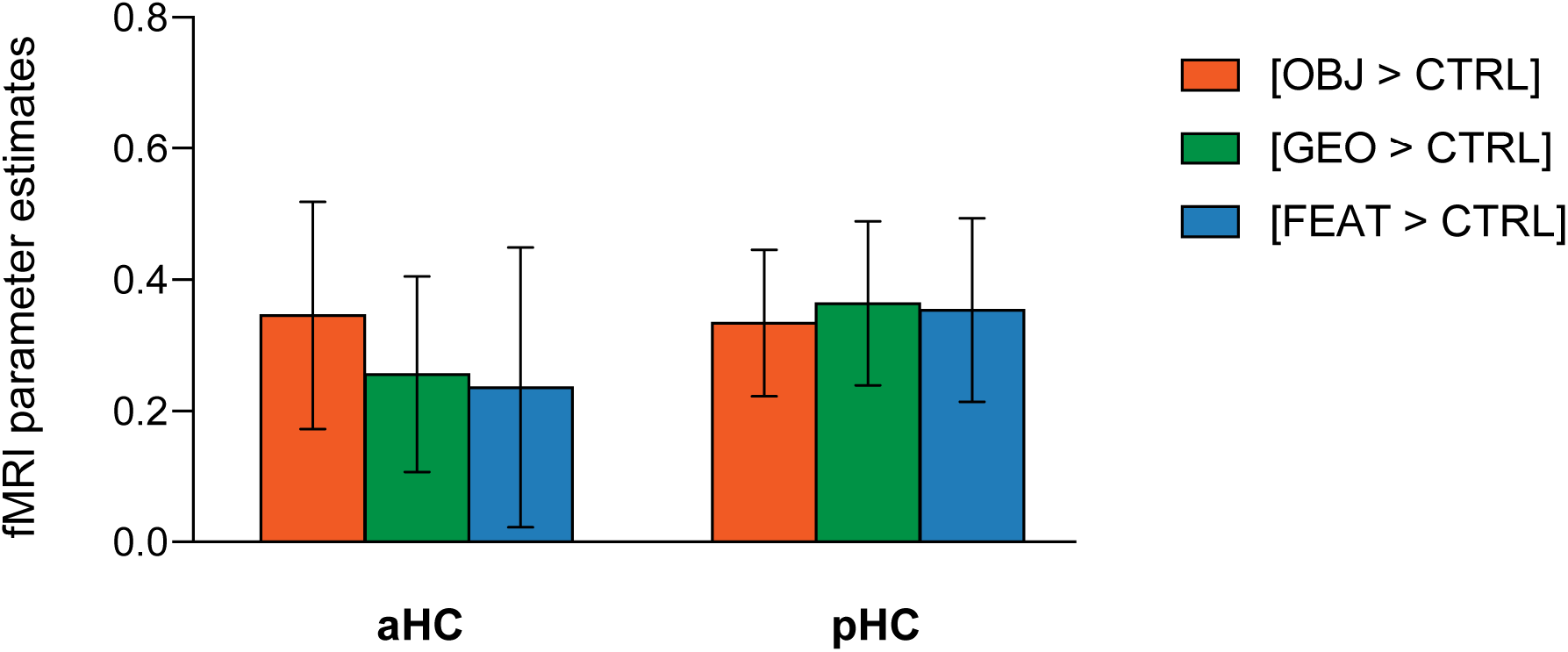
fMRI parameter estimates in the anterior hippocampus (aHC) and posterior hippocampus (pHC) for the fMRI contrasts [OBJ > CTRL], [GEO > CTRL] and [FEAT > CTRL]. Error bars reflect standard errors of the mean.

**Table S1.** Descriptive characteristics and cognitive performance of participants. M: male; F: female; SEM: standard error of the mean; MMSE: mini mental state examination.

| Variables | Mean (± SEM) |
| --- | --- |
| Age | 25.4 (± 0.5) |
| Males / Females | 18 / 7 |
| Total brain volume (cm<sup>3</sup>) | 1301 (± 18) |
| MMSE | 30.0 (± 0.0) |
| 3D mental rotation | 18.3 (± 0.9) |
| Corsi forward | 7.2 (± 0.2) |
| Corsi backward | 6.2 (± 0.3) |
| Perspective taking test | 15.3 (± 1.7) |

**Table S2.** Cerebral regions whose activity was elicited by the object condition in comparison with the control condition (and reciprocally). The statistical threshold was defined as *p* < 0.001 uncorrected for multiple comparisons with an extent voxel threshold defined as 10 voxels. For each cluster, the region showing the maximum t-value was listed first, followed by the other regions in the cluster [in square brackets]. Montreal Neurological Institute (MNI) coordinates (x, y, z) of the peak and number of voxels (k) of clusters are also shown. OBJ = object condition; CTRL = control condition; H = hemisphere; R = right; L = left; BA = Brodmann area; CI = confidence interval.

|  |  | H | BA | k | x | y | z | t | ES [95%-CI] |
| --- | --- | --- | --- | --- | --- | --- | --- | --- | --- |
| Contrasts one-sample |  |  |  |  |  |  |  |  |  |
| [OBJ > CTRL] | Cerebellum (Crus I) | R |  | 365 | 21 | -61 | -34 | 6.55 | 0.95 [0.66-1.23] |
|  | [Cerebellum Lobule VI] |  |  |  | 18 | -70 | -28 | 5.22 | 1.17 [0.73-1.61] |
|  | [Middle Occipital Gyrus] |  |  |  | 18 | -100 | -1 | 5.08 | 3.08 [1.89-4.26] |
|  | [Middle Occipital Gyrus] |  |  |  | 42 | -82 | -4 | 4.10 | 1.58 [0.95-2.22] |
|  | Inferior Temporal Gyrus | R | 38 | 23 | 36 | 8 | -34 | 5.84 | 0.85 [0.56-1.14] |
|  | [Middle Temporal Gyrus] |  | 20 |  | 45 | 8 | -37 | 5.03 | 1.02 [0.62-1.42] |
|  | Fusiform Gyrus | L | 37 | 86 | -36 | -52 | -16 | 5.50 | 1.44 [0.93-1.96] |
|  | [Inferior Temporal Gyrus] |  |  |  | -51 | -58 | -19 | 3.73 | 1.71 [0.81-2.61] |
|  | Angular Gyrus | R | 39 | 34 | 36 | -55 | 17 | 5.32 | 0.79 [0.50-1.08] |
|  | Middle Occipital Gyrus | L | 19 | 154 | -42 | -85 | -10 | 5.14 | 1.98 [1.23-2.74] |
|  | [Middle Occipital Gyrus] |  | 18 |  | -24 | -100 | -1 | 5.09 | 2.66 [1.63-3.68] |
|  | [Inferior Occipital Gyrus] |  | 18 |  | -27 | -94 | -10 | 4.84 | 2.27 [1.35-3.19] |
|  | Hippocampus | L | 54 | 35 | -33 | -22 | -13 | 5.07 | 0.98 [0.60-1.36] |
|  | [Hippocampus] |  | 54 |  | -30 | -10 | -16 | 4.49 | 0.95 [0.54-1.36] |
|  | [Sup. Temporal Gyrus] |  | 38 |  | -33 | 5 | -22 | 3.76 | 1.79 [0.86-2.72] |
|  | Anterior Cingulate Gyrus | R | 32 | 35 | 6 | 38 | 5 | 4.91 | 0.94 [0.60-1.36] |
|  | Inferior Frontal Gyrus | L | 44 | 11 | -60 | 14 | 26 | 4.66 | 1.10 [0.63-1.56] |
|  | Cerebellum (Lobule VIIb) | R |  | 26 | 21 | -73 | -49 | 4.66 | 1.16 [0.67-1.66] |
|  | [Cerebellum Crus II] |  |  |  | 15 | -79 | -40 | 4.62 | 1.02 [0.56-1.48] |
|  | Inferior Occipital Gyrus | R | 19 | 75 | 39 | -64 | -7 | 4.49 | 1.38 [0.78-1.98] |
|  | [Fusiform Gyrus] |  | 37 |  | 42 | -58 | -13 | 4.39 | 1.72 [0.95-2.49] |
|  | [Inferior Temporal Gyrus] |  | 37 |  | 51 | -55 | -22 | 4.09 | 2.29 [1.19-3.39] |
|  | Lat. Orbitofrontal Gyrus | L | 47 | 18 | -48 | 23 | -13 | 4.43 | 1.99 [1.11-2.88] |
|  | Superior Temporal Gyrus | L | 22 | 10 | -63 | -25 | 5 | 4.35 | 1.23 [0.67-1.78] |
|  | Superior Temporal Gyrus | R | 22 | 19 | 60 | -25 | 2 | 4.29 | 1.57 [0.85-2.28] |
| [CTRL > OBJ] |  | No significant activation |  |  |  |  |  |  |  |

**Table S3.** Cerebral regions whose activity was elicited by the geometry-based condition in comparison with the control condition (and reciprocally). The statistical threshold was defined as *p* < 0.001 uncorrected for multiple comparisons with an extent voxel threshold defined as 10 voxels. For each cluster, the region showing the maximum t-value was listed first, followed by the other regions in the cluster [in square brackets]. Montreal Neurological Institute (MNI) coordinates (x, y, z) of the peak and number of voxels (k) of clusters are also shown. GEO = geometry; CTRL = control; H = hemisphere; R = right; L = left; BA = Brodmann area, CI = confidence interval.

|  |  | H | BA | k | x | y | z | t | ES [95%-CI] |
| --- | --- | --- | --- | --- | --- | --- | --- | --- | --- |
| **Contrasts one-sample** |  |  |  |  |  |  |  |  |  |
| **[GEO > CTRL]** | Anterior Cingulate Gyrus | L/R | 24 | 122 | 6 | 32 | 2 | 6.02 | 1.25 [0.84-1.65] |
|  | [Superior Frontal Gyrus] |  |  |  | 3 | 35 | 26 | 4.43 | 1.80 [1.00-2.60] |
|  | [Middle Occipital Gyrus] |  |  |  | 18 | -100 | -1 | 5.08 | 1.69 [0.94-2.44] |
|  | Inferior Temporal Gyrus | R | 37 | 80 | 42 | -58 | -7 | 5.64 | 1.72 [1.12-2.32] |
|  | [Middle Temporal Gyrus] |  | 19 |  | 54 | -64 | 5 | 4.62 | 2.03 [1.17-2.89] |
|  | [Angular Gyrus] |  | 37 |  | 42 | -52 | 11 | 3.90 | 1.31 [0.65-1.96] |
|  | Cerebellum (Lob. I-IV) | L/R |  | 24 | 0 | -49 | -19 | 5.38 | 1.48 [0.94-2.02] |
|  | Superior Frontal Gyrus | L/R | 10 | 112 | 3 | 50 | 8 | 5.29 | 2.40 [1.51-3.28] |
|  | [] |  |  |  | 9 | 56 | 14 | 5.11 | 1.21 [0.75-1.68] |
|  | Middle Temporal Gyrus | L | 19 | 57 | -54 | -67 | -4 | 5.07 | 2.30 [1.41-3.20] |
|  | Inferior Occipital Gyrus | L | 18 | 42 | -27 | -91 | -10 | 4.55 | 2.70 [1.54-3.86] |
|  | Cerebellum | R |  | 10 | 18 | -61 | -34 | 4.48 | 0.72 [0.40-1.03] |
|  | Thalamus | R |  | 16 | 27 | -25 | 14 | 4.40 | 0.90 [0.50-1.30] |
|  | [Caudate] |  |  |  | 21 | -16 | 20 | 3.79 | 0.93 [0.45-1.41] |
|  | Cerebellum (Lob. IX) | R |  | 32 | 9 | -52 | -40 | 4.38 | 1.33 [0.74-1.93] |
|  | [Cerebellum Vermis VIII] |  |  |  | 6 | -64 | -37 | 4.05 | 1.02 [0.52-1.51] |
|  | Parahippocampal Gyrus | L | 36 | 25 | -30 | -37 | -13 | 4.38 | 1.20 [0.66-1.74] |
|  | Inferior Occipital Gyrus | R | 18 | 39 | 33 | -85 | -7 | 4.31 | 2.70 [1.47-3.93] |
|  | Precentral Gyrus | L | 6 | 26 | -30 | -10 | 65 | 4.27 | 1.73 [0.94-2.53] |
|  | [Superior Frontal Gyrus] |  |  |  | -21 | -7 | 65 | 4.03 | 1.16 [0.60-1.72] |
|  | Insula | L | 13 | 19 | -36 | -1 | 5 | 4.27 | 1.11 [0.60-1.62] |
|  | Sup. Temporal Gyrus | L | 22 | 10 | -57 | 2 | -10 | 4.11 | 1.16 [0.61-1.71] |
|  | Middle Temporal Gyrus | L | 21 | 14 | -63 | -16 | -16 | 4.02 | 1.03 [0.53-1.53] |
|  | Fusiform Gyrus | R | 36 | 19 | 33 | -43 | -7 | 3.91 | 0.90 [0.45-1.36] |
| **[CTRL > GEO]** |  | *No significant activation* |  |  |  |  |  |  |  |

**Table S4.** Cerebral regions whose activity was elicited by the feature-based condition in comparison with the control condition (and reciprocally). The statistical threshold was defined as *p* < 0.001 uncorrected for multiple comparisons with an extent voxel threshold defined as 10 voxels. For each cluster, the region showing the maximum t-value was listed first, followed by the other regions in the cluster [in square brackets]. Montreal Neurological Institute (MNI) coordinates (x, y, z) of the peak and number of voxels (k) of clusters are also shown. FEAT = feature; CTRL = control; H = hemisphere; R = right; L = left; BA = Brodmann area; CI = confidence interval.

|  |  | H | BA | k | x | y | z | t | ES [95%-CI] |
| --- | --- | --- | --- | --- | --- | --- | --- | --- | --- |
| **Contrasts one-sample** |  |  |  |  |  |  |  |  |  |
| **[FEAT > CTRL]** | Inferior Frontal Gyrus | L | 44 | 16 | -60 | 14 | 26 | 5.26 | 1.19 [0.74-1.63] |
|  | [Precentral Gyrus] |  | 6 |  | -63 | 2 | 29 | 3.28 | 1.39 [0.66-2.12] |
|  | Superior Frontal Gyrus | R | 11 | 14 | 0 | 38 | -19 | 5.13 | 1.19 [0.73-1.64] |
|  | Inferior Occipital Gyrus | R | 19 | 30 | 39 | -64 | -7 | 5.02 | 1.19 [0.73-1.66] |
|  | Fusiform Gyrus | R | 37 | 28 | 33 | -31 | -19 | 4.70 | 1.07 [0.62-1.51] |
|  | Anterior Cingulate Gyrus | R |  | 46 | 9 | 32 | 5 | 4.52 | 1.08 [0.61-1.55] |
|  | □ |  |  |  | -6 | 26 | 14 | 3.61 | 0.66 [0.30-1.02] |
|  | Cerebellum | R |  | 17 | 27 | -55 | -43 | 4.45 | 0.91 [0.51-1.31] |
|  | Inferior Occipital Gyrus | R | 19 | 13 | 30 | -82 | -13 | 4.33 | 2.81 [1.54-4.08] |
|  | Postcentral Gyrus | L | 4 | 10 | -21 | -28 | 68 | 4.26 | 1.09 [0.59-1.60] |
|  | Hippocampus | L | 54 | 10 | -33 | -25 | -13 | 4.24 | 0.87 [0.47-1.28] |
| **[CTRL > FEAT] No significant activation** |  |  |  |  |  |  |  |  |  |

**Table S5.** Cerebral regions whose activity was elicited by direct comparison between the cue-based conditions. The statistical threshold was defined as *p* < 0.001 uncorrected for multiple comparisons with an extent voxel threshold defined as 10 voxels. For each cluster, the region showing the maximum t-value was listed first, followed by the other regions in the cluster [in square brackets]. Montreal Neurological Institute (MNI) coordinates (x, y, z) of the peak and number of voxels (k) of clusters are also shown. OBJ = object condition; GEO: geometry condition; FEAT: feature condition; H = hemisphere; R = right; L = left; BA = Brodmann area; CI = confidence interval.

|  |  | H | BA | k | x | y | z | t | ES [95%-CI] |
| --- | --- | --- | --- | --- | --- | --- | --- | --- | --- |
| **Contrasts one-sample** |  |  |  |  |  |  |  |  |  |
| [OBJ > GEO] | *No significant activation* |  |  |  |  |  |  |  |  |
| [GEO > OBJ] | Superior Frontal Gyrus | L | 6 | 52 | -18 | -4 | 65 | 5.08 | 9.94 [6.10-13.77] |
|  | □ |  |  |  | -27 | -4 | 74 | 4.52 | 11.87 [6.72-17.02] |
|  | [Precentral Gyrus] |  |  |  | -30 | -10 | 62 | 3.72 | 9.78 [4.63-14.94] |
|  | Brainstem | L/R |  | 11 | 0 | -22 | -16 | 5.02 | 8.13 [4.96-11.20] |
|  | Caudate Nucleus | L | 48 | 10 | -9 | 11 | -1 | 4.76 | 9.04 [5.32-12.76] |
|  | Superior Frontal Gyrus | L | 6 | 16 | -9 | -16 | 62 | 4.56 | 5.93 [3.38-8.47] |
|  | [Precentral Gyrus] |  |  |  | -3 | -16 | 56 | 4.40 | 8.90 [4.93-12.88] |
|  | Cerebellum | L |  | 12 | -12 | -55 | -43 | 4.27 | 5.43 [2.94-7.92] |
|  | Caudate Nucleus | L | 48 | 17 | -12 | 2 | 11 | 4.06 | 11.29 [5.83-16.74] |
|  | □ |  |  |  | -15 | 2 | 20 | 3.62 | 9.27 [4.25-14.30] |
| [OBJ > FEAT] | *No significant activation* |  |  |  |  |  |  |  |  |
| [FEAT > OBJ] | *No significant activation* |  |  |  |  |  |  |  |  |
| [GEO > FEAT] | Middle Temporal Gyrus | L | 37 | 38 | -54 | -64 | -1 | 5.48 | 12.22 [7.84-16.60] |
|  | Middle Frontal Gyrus | R | 9 | 21 | 30 | 44 | 35 | 5.20 | 9.06 [5.65-12.48] |
|  | [Superior Frontal Gyrus] |  |  |  | 24 | 35 | 32 | 4.17 | 4.41 [2.34-6.48] |
|  | Inferior Occipital Gyrus | L | 18 | 44 | -27 | -88 | -10 | 5.13 | 11.95 [7.38-16.51] |
|  | Middle Temporal Gyrus | R | 37 | 19 | 51 | -61 | -1 | 4.91 | 9.97 [5.99-13.95] |
|  | Putamen | R | 49 | 17 | 33 | -16 | 2 | 4.70 | 7.12 [4.15-10.09] |
|  | Superior Parietal Gyrus | L | 7 | 21 | -21 | -52 | 65 | 4.39 | 7.89 [4.36-11.41] |
|  | Putamen | L | 49 | 14 | -30 | 5 | -1 | 4.07 | 4.67 [2.42-6.91] |
|  | Middle Frontal Gyrus | L | 10 | 11 | -27 | 47 | 26 | 4.06 | 8.44 [4.37-12.52] |
|  | Supramarginal Gyrus | R | 40 | 16 | 51 | -31 | 35 | 3.89 | 8.77 [4.35-13.19] |
| [FEAT > GEO] | *No significant activation* |  |  |  |  |  |  |  |  |

## Notes

### Competing Interest Statement

The authors have declared no competing interest.

### Summary of Updates

Representational similarity analyses were added to the Methods and Results sections. Figure 5 was revised accordingly.

